# Correlating cell function and morphology by performing fluorescent immunocytochemical staining on the light-microscope stage

**DOI:** 10.1101/2020.06.30.180810

**Authors:** Hiroyuki Kawano, Yasuhiro Kakazu, Sadahiro Iwabuchi, N. Charles Harata

**Affiliations:** Department of Molecular Physiology & Biophysics, University of Iowa Carver College of Medicine, Iowa City, Iowa, United States of America; Kakazu ENT Clinic, Higashi-ku, Fukuoka, Japan; Department of Molecular Pathophysiology, Institute of Advanced Medicine, Wakayama Medical University, Wakayama, Wakayama, Japan; Iowa Neuroscience Institute, University of Iowa Carver College of Medicine, Iowa City, Iowa, United States of America

**Author notes:** Corresponding author (NCH).

**Keywords:** Calcium concentration, Chemical fixation, Fluorescence imaging, Image translation, Immunocytochemistry, Presynaptically silent synapses

## Abstract

**Background:** Correlation of fluorescence signals from functional changes in live cells with those from immunocytochemical indicators of their morphology following chemical fixation can be highly informative with regard to function-structure relationship. Such analyses can be technically challenging because they need consistently aligning the images between imaging sessions. Existing solutions include introducing artificial spatial landmarks and modifying the microscopes. However, these methods can require extensive changes to the experimental systems.

**New method:** Here we introduce a simple approach for aligning images. It is based on two procedures: performing immunocytochemistry while a specimen stays on a microscope stage (on-stage), and aligning images using biological structures as landmarks after they are observed with transmitted-light optics in combination with fluorescence-filter sets.

**Results:** We imaged a transient functional signal from a fluorescent Ca^2+^ indicator, and mapped it to neurites based on immunocytochemical staining of a structural marker. In the same preparation, we could identify presynaptically silent synapses, based on a lack of labeling with an indicator for synaptic vesicle recycling and on positive immunocytochemical staining for a structural marker of nerve terminals. On-stage immunocytochemistry minimized lateral translations and eliminated rotations, and transmitted-light images of neurites were sufficiently clear to enable spatial registration, effective at a single-pixel level.

**Comparison with existing methods:** This method aligned images with minimal change or investment in the experimental systems.

**Conclusions:** This method facilitates information retrieval across multiple imaging sessions, even when functional signals are transient or local, and when fluorescent signals in multiple imaging sessions do not match spatially.

## 1. Introduction

Fluorescence, the light emitted by fluorescent probes (fluorophores), is commonly used in the imaging of biological cells and tissues at the light microscope level (Giepmans et al., 2006; Pawley, 2006; Rodriguez et al., 2017). Changes in either the intensity or lifetime of fluorescence carry information about the changes in fluorophore microenvironment. These changes are used for effectively monitoring functions of live cells, such as regulations of cytosolic Ca^2+^ concentration ([Ca^2+^]_c_) (Dana et al., 2019; Iwabuchi et al., 2013b) and membrane potential (Fan et al., 2020; Knopfel and Song, 2019; Miller et al., 2012). In addition, the location and intensity of fluorescence are sensitive indicators of the location and amount of a fluorophore. These indications are used for effectively collecting morphological information in the fluorescence-based immunocytochemistry (ICC), where cells are chemically fixed and the fluorophore-conjugated antibodies are used to detect an intracellular antigen of interest (Kawano et al., 2012; Mitchell et al., 2018; Schneider Gasser et al., 2006) (as opposed to extracellular antigens which do not require chemical fixations) (**Fig. 1A**).

**Fig. 1.**
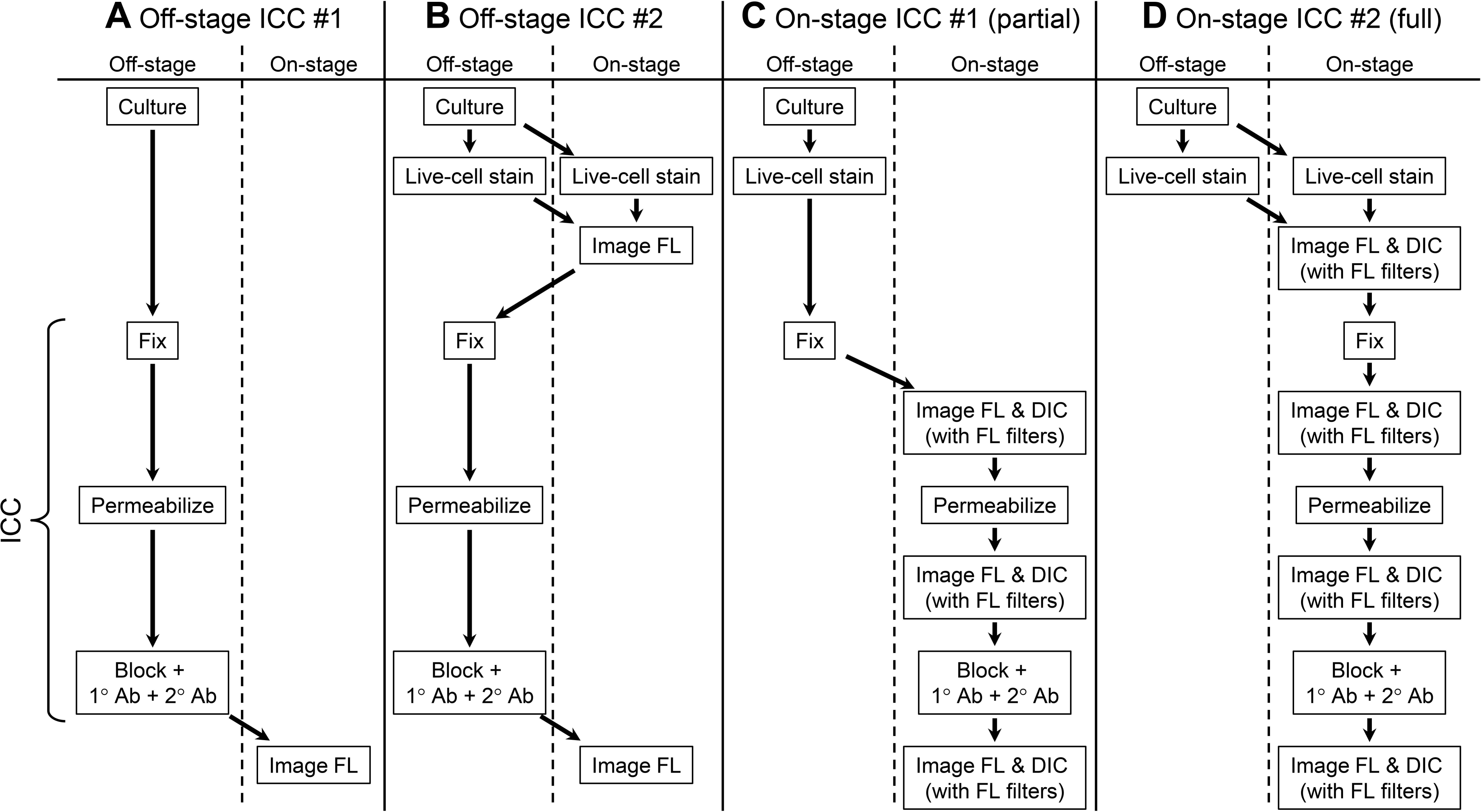
Flowcharts of procedures for fixed-cell immunocytochemistry (ICC). The listed steps are performed either outside (off-stage) or on the light-microscope stage (on-stage), as indicated. (**A**) Classical ICC protocol that does not involve live-cell fluorescence imaging (Off-stage ICC #1). A curly bracket on left indicates the ICC procedures from chemical fixation to treatments with primary and secondary antibodies, which are applicable to all four flowcharts (**A-D**). (**B**) Standard ICC protocol that is preceded by live-cell fluorescence imaging (Off-stage ICC #2). (**C**) One version of our method (On-stage ICC #1; partial): Procedures up to chemical fixation are performed off-stage and then all procedures are performed on-stage. (**D**) Another version of our method (On-stage ICC #2; full): All key procedures are performed on-stage. In **C** and **D**, live-cell staining can be done off- or on-stage. **Abbreviations** 1° Ab: primary antibody; 2° Ab: secondary antibody conjugated with fluorophores; Block: blocking of non-specific binding of antibodies; DIC: differential interference contrast transmitted-light microscopy; Fix: chemical fixation, e.g. using aldehyde-based fixatives; FL: fluorescence microscopy; FL filters: fluorescence filters; ICC: immunocytochemistry, based on immuno-fluorescence method; Stage: microscope stage, which holds the imaging chamber and biological specimen.

Combining the imaging information of the same specimens acquired under both live- and fixed-cell conditions is expected to yield deep insight into cellular functions and structures. Typical experiments for combining the information proceed as follows (**Fig. 1B**). A live specimen, e.g. cultured cells on a glass coverslip, is placed in an imaging chamber, which in turn is placed onto a light-microscope stage (on-stage) through an adapter. Images of the live cells are acquired using a camera attached to the microscope body. Then the specimen will be brought to a laboratory bench (off-stage), be chemically fixed in preparation for ICC of intracellular antigens, and the fixed cells will be stained with antibodies against the structural antigens of interest. After the specimen is returned to the microscope stage, the field imaged during the live-cell session is reimaged. Post-acquisition analysis starts with alignment of the images acquired under live- and fixed-cell conditions, via lateral translation in the x-y dimension and rotation around the z-axis, such that the fluorescence signals should coincide spatially.

However, when live- and fixed-cell imaging is combined in this way, there can arise at least five problems that are associated with spatial and temporal mismatches. Two are procedural (1 and 2 below), two are biological (3 and 4), and one (5) is optical. Problem 1 is that, between live- and fixed-cell imaging sessions, the position of a specimen can shift with respect to the camera. This can occur either because the microscope stage is moved with respect to the microscope body or because the specimen or imaging chamber moves with respect to the microscope stage even when the stage is held stationary. This type of movement is almost inevitable, especially if the rotation angle of the imaging chamber can change freely with respect to the microscope stage. Due to Problem 1, the specimen will appear to have shifted laterally and/or rotated between the first and second sessions. Of the five problems, this one accounts for the most sample movement.

Problem 2 is that misalignment of optical components of a microscope can lead to image movements. Typically such misalignment is caused by mechanical switching of fluorescence filter cubes, and/or by switching among different fluorescence-excitation light sources (when multiple fluorescence channels are used). Typical modes of transmitted light microscopy, such as differential interference contrast (DIC) microscopy and phase-contrast microscopy, do not require a fluorescence filter cube, whereas the fluorescence imaging requires at least one. Thus, as filter cubes are moved mechanically in a single experiment, they can cause slight changes in the light path, and thereby the lateral shift of images.

Problem 3 is that spatial locations differ by the nature of signal sources in two imaging sessions. An example is found in the study of presynaptically silent synapses. A certain proportion of nerve terminals do not release neurotransmitter even when neurons are stimulated extensively (Crawford and Mennerick, 2012; Kawano et al., 2012; Moulder et al., 2004): these presynaptically silent synapses do not take up FM dye, a fluorescent marker of the synaptic-vesicle recycling (Iwabuchi et al., 2014a; Kavalali and Jorgensen, 2014; Kawano et al., 2012), although those nerve terminals are morphologically mature. Thus, such nerve terminals lack fluorescent FM signals yet are positive for ICC signals for structural markers of nerve terminals. In this case, fluorescence signals obtained during one imaging session will not necessarily colocalize spatially with those in another. Thus, fluorescent signals are not necessarily a reliable means of identifying specimen movement, even when they are emitted from the same specimen.

Problem 4 is that spatio-temporal relationships can differ from one imaging session to another. For example, functional changes in [Ca^2+^]_c_ are transient, with [Ca^2+^]_c_ waves propagating through the cytosol (Ross, 2012). In addition, they can be confined in space, as in [Ca^2+^]_c_ sparks which represent local and discrete elevations in [Ca^2+^]_c_ (Brochet et al., 2012; Cheng and Lederer, 2008; Ross, 2012). These functional signals are absent in fixed cells/tissues, and their spatial domains do not necessarily match morphological spaces, e.g. those occupied by the dendrites and somata of neurons. Thus, the live-cell signal will not provide a spatial or morphological cue during the fixed-cell imaging step.

Problem 5 is that spatial displacement can arise from lateral chromatic aberrations. The projection of light by optical components of a microscope varies by wavelength (Hibbs et al., 2006). This leads to lateral displacement of images at different wavelengths, even when the signal is emitted from a multi-color fluorophore. Such displacement creates the smallest amount of specimen movement among the five problems.

Although approaches to solving these problems exist, they all have limitations. For Problem 1, one approach is to modify the protocol: ICC procedures are performed off-stage but with the specimen still in place within the imaging chamber. If the imaging chamber is returned to the microscope stage with care, movements between imaging sessions will be reduced. But some movement will still remain and be detectable at the cellular level. For Problem 1, another approach is to locate and align fields of interest during imaging sessions, using points of spatial reference, i.e. non-biological landmarks (fiducial marks/markers) that are visible during live- and fixed-cell imaging sessions. Exemplar methods for locating imaged fields include the use of glass coverslips etched with a specific grid pattern (non-fluorescent but visible with transmitted light; e.g. CELLocate, Eppendorf), the use of coverslips printed with gold nanoparticles (Prabhakar et al., 2018), and near-infrared branding of specimens with artificial autofluorescence (Bishop et al., 2011). After the imaged fields are located during the second session, they are aligned to the previously acquired images by manually translating the microscope stage in the x-y dimension and rotating the imaging chamber around the optical z-axis, e.g. using an adaptor (specimen holder) with a rotation control (Iwabuchi et al., 2014c). However, this solution is effective only if the landmarks are labeled at a density sufficient to be visible during high-magnification imaging of subcellular structures. This is often challenging, with the landmarks remaining outside of the target field.

For Problems 1 and 2, a general approach to solving mismatch is to align the acquired fluorescence images manually using image-analysis software, e.g. NIH ImageJ and Adobe Photoshop. The image movements can be corrected by optimizing overlap of fluorescence signals from two or more channels. However, this post-acquisition approach is not effective when fiducial markers are applied at too low a density to be visible in the imaged field, or when biological fluorescent signals are not present at the same spatial location in multiple image sessions as in Problems 3 and 4. Correcting movements by image rotation also has the limitation that the fluorescence intensity in the original pixel is spread among multiple neighboring pixels by the interpolation transformation built into image-analysis software (p. 261 in Russ, 2011). Under standard imaging conditions, e.g. by widefield optical microscopy, this is a serious problem, especially for target structures that provide punctate signals of only at most a few pixels wide, such as nerve terminals.

Here we report on a simple approach for minimizing spatial and temporal mismatch between signals under live-vs. fixed-cell conditions. It is based on two procedures and is intended for imaging subcellular structures using a light microscope at a micrometer or pixel level. First, ICC procedures are performed on-stage to suppress unintentional movement of the specimen with respect to the camera and to limit any potential residual displacement to a slight lateral translation in the x-y plane with no rotation. Second, residual displacement is corrected after image acquisition using biological structures (e.g. neurites) as landmarks. Those structures are imaged using transmitted-light optics (DIC or phase-contrast microscopy) in combination with fluorescence-filter sets when the fluorescence images are acquired. After the images are acquired, the DIC images are aligned and the same changes in coordinates (transformation or translation) are used to align fluorescence images.

Our approach has the advantage that it collectively corrects all lateral displacements, irrespective of the causes. It is also simple, yet capable of solving most of the problems that lead to mismatch. The sole exception is chromatic aberration (Problem 5), which has a minimal effect. We describe the principle underlying the on-stage ICC procedures, as well as two versions of this approach (**Figs. 1C**,**D**). We also illustrate its usefulness in experiments where [Ca^2+^]_c_ transients and presynaptically silent synapses are observed in cultured mouse hippocampal neurons.

## 2. Materials and Methods

### 2.1. Ethics statement

Animal care and use procedures were approved by the University of Iowa’s Institutional Animal Care and Use Committee, and performed in accordance with the standards set by the PHS Policy and The Guide for the Care and Use of Laboratory Animals (NRC Publications) revised 2011. Every effort was made to minimize suffering of the animals.

### 2.2. Cell culture

Neurons in primary culture were prepared from wild-type mice of both sexes on postnatal days 0-1, by methods described previously (Koh et al., 2015; Mitchell et al., 2018) with slight modifications. The CA3-CA1 region of the hippocampus was dissected bilaterally and treated with trypsin (from bovine pancreas, 2.5 mg/ml, T1005; MilliporeSigma, Burlington, Massachusetts, USA) and deoxyribonuclease I (DNase from bovine pancreas, 750 units/ml, D4527; MilliporeSigma) for 13 min at room temperature, and mechanically dissociated. The cells were centrifuged for 13 min at ∼185 g at 4°C. The pellet was suspended, and multiple neurons were plated on 12-mm coverslips (thickness No. 0; Carolina Biological Supply, Burlington, North Carolina, USA) in 24-well plates, at a density of 10,000 cells per coverslip (mass culture). The coverslips had previously been seeded with an astrocytic feeder layer at 10,000 or 20,000 cells per coverslip for 7-9 days. Feeder cells were prepared from the CA3-CA1 region of the wild-type mouse hippocampus on postnatal days 0-1. The neurons were cultured in a humidified incubator at 37°C with 5% CO_2_ until they were used for experiments at 13-16 days *in vitro*.

### 2.3. Live-cell staining of functional nerve terminals with FM dye for partial on-stage ICC

The procedures for the partial on-stage ICC (**Fig. 1C**) are described in sections 2.3-2.5. The additional procedures for the full on-stage ICC (**Fig. 1D**) are described in sections 2.6-2.9. Some detailed comments on the procedures are listed in the Supplementary Information.

Functional nerve terminals (boutons) were stained by labeling recycling synaptic vesicles with FM1-43 (T35356; Thermo Fisher Scientific, Waltham, Massachusetts, USA) under a condition promoting high neuronal activity (Iwabuchi et al., 2014a; Kakazu et al., 2012a), off-stage, at room temperature (∼21°C). The coverslips were immersed in 10 μM FM1-43 in a high-K^+^ solution (45-mM KCl) for 1 min. The high-K^+^ extracellular solution was prepared by equimolar substitution of KCl for NaCl. The dye was washed out for 5 min, by transferal of the coverslip into a culture dish filled with dye-free Tyrode’s solution containing 1 μM of the voltage-gated Na^+^ channel blocker tetrodotoxin (tetrodotoxin citrate, TTX, 1069; Tocris Bioscience, Minneapolis, Minnesota, USA) to block loss of the stained FM dye due to the firing of spontaneous action potentials. The stock solutions of FM1-43 and TTX were prepared at 2.5 mM and 1 mM, respectively, in distilled water, and stored at -80°C.

### 2.4. Fixed-cell staining of neurons during partial on-stage ICC

ICC was used to localize: a glutamatergic presynaptic marker, vesicular glutamate transporter 1 (VGLUT1); a GABAergic presynaptic marker, vesicular GABA transporter (VGAT); and a dendritic marker, microtubule-associated protein 2 (MAP2). Basic features of the ICC procedures were generally the same as in previous reports (Moulder et al., 2008; Moulder et al., 2010), but some details were modified, as specified in the Supplementary Information. All procedures were performed at room temperature.

Immediately after being stained with the FM dye, the cultured neurons were fixed for 10-15 min by transferal of the coverslip into a dish with 4% paraformaldehyde (16% aqueous solution, 15710; Electron Microscopy Sciences, Hatfield, Pennsylvania, USA) and 0.2% glutaraldehyde (25% aqueous solution, 16220; Electron Microscopy Sciences) (Moulder et al., 2010). The solvent was phosphate-buffered saline (PBS, 10× that was diluted to 1×, 70011-044; Thermo Fisher Scientific) adjusted to pH 7.4 using 0.037% hydrochloric acid (258148; MilliporeSigma) and its osmolarity was determined to be ∼299 mOsm without any addition of sucrose. After being rinsed by transferal into fresh PBS twice, 5 min each, coverslips were placed in an open-bath imaging chamber (RC-26; Warner Instruments, Hamden, Connecticut, USA). The floor of the imaging chamber was a rectangular coverslip (22 × 40 mm, thickness 0.08-0.13 mm, No. 0, 72198-20; Electron Microscopy Sciences). The chamber was fitted onto a platform with solution inlets (PH-1, MCK-1; Warner Instruments) and a stage adaptor (SA-NIK; Warner Instruments), and then onto the stage of an inverted widefield fluorescence microscope (Eclipse Ti-E; Nikon, Melville, New York, USA). The chamber volume was set to ∼700 μl by adjusting the height of a suction tube. All of the following procedures were performed on-stage. The coverslip was continuously perfused (using the bath-perfusion line through a gravity-driven drip-infusion system) with PBS at a rate of ∼20 μl/sec for ∼30 min (total volume of PBS ∼36 ml). During this washing period, images were captured, using DIC optics for cell morphology and fluorescence optics for the FM dye. Then the cells were permeabilized and blocked for 15 min with 0.1% Triton X-100 (T8787; MilliporeSigma) and 4% normal goat serum (NGS, 005-000-121; Jackson ImmunoResearch Laboratories, West Grove, Pennsylvania, USA) in PBS. The volume of this solution totaled 5-10 ml and it was applied through the first of the two inlet lines, which were separate from the bath-perfusion line, while the bath perfusion was stopped. Subsequently, the cells were washed with 4% NGS in PBS (final concentration, blocking solution) through the first inlet line for 15 min (total volume of 30-40 ml); they were simultaneously washed with PBS through the bath-perfusion line. During the last few minutes of washing, the bath perfusion was stopped to keep the NGS concentration in the bath at ∼4%.

The cells were then treated for 3 hours with a primary antibody solution containing polyclonal guinea-pig anti-VGLUT1 antibody (AB5905; MilliporeSigma) (2,000× dilution), monoclonal mouse anti-VGAT antibody (131 011; Synaptic Systems, Göttingen, Germany) (2,000× dilution), and polyclonal rabbit anti-MAP2 antibody (AB5622; MilliporeSigma) (2,000× dilution) in blocking solution. The total volume of the primary antibody solution was 2 ml and it was applied through the first inlet line while the bath perfusion was stopped. Subsequently, the cells were washed, for 15 min, with blocking solution applied via the first inlet line (30-40 ml) and simultaneously with PBS applied via the bath perfusion line. The cells were then treated for 60-90 min with a secondary antibody solution containing goat anti-guinea-pig IgG (H+L) antibody conjugated with Alexa Fluor 594 (A-11076; Thermo Fisher Scientific) (1,000× dilution), goat anti-mouse IgG (H+L) antibody conjugated with Alexa Fluor 488 (A-11001; Thermo Fisher Scientific) (1,000× dilution), and goat anti-rabbit IgG (H+L) antibody conjugated with Alexa Fluor 405 (A-31556; Thermo Fisher Scientific) (1,000× dilution) in blocking solution. The total volume of the secondary antibody solution was 2 ml, and it was applied through a second inlet line while the bath perfusion was stopped. The cells were then washed with PBS through the second inlet line (30-40 ml) for 30 min, and simultaneously with PBS through the bath perfusion line. Each of the two inlet lines had an internal volume (i.e. a dead volume) of 800-900 μl. Images of neurons stained by ICC were captured while the coverslips were immersed in PBS.

### 2.5. Fluorescence imaging of neurons stained with FM dye and with ICC during partial on-stage ICC

After neurons were stained with FM dye, those to be imaged were selected using DIC optics as a guide, without prior information about fluorescence. This procedure avoided biased acquisition of fluorescence images during subsequent steps. The selection was based on the presence of features of healthy neurons, including: a clear cellular margin; extended, non-beaded dendrites; a uniform glial layer; lack of clustering of multiple somata; and lack of bundling of neuronal processes. Once the neurons were selected, their 14-bit DIC and fluorescence images were acquired, using an interline charge-coupled device (CCD) camera (Clara; Andor Technology, Springvale Business Park, Belfast, UK) and the following parameters: full-frame format without binning; pre-amplifier gain 1×; extended near-infrared (NIR) mode ON; and at 10-MHz pixel readout rate. The camera was cooled at -45°C by an internal fan to reduce camera noise. Cells were imaged using an inverted microscope (Eclipse Ti-E; Nikon), with a 40× oil-immersion objective lens (numerical aperture 1.30, Plan Fluor; Nikon), without an intermediate coupler (i.e. 1×), and with an exposure time of 50 ms for DIC imaging and 1 s for fluorescence imaging. FM1-43 was excited using a 490-nm light-emitting diode (LED, precisExcite LED array module; CoolLED, Andover, Hampshire, UK) at 100% intensity, and a “green” filter cube (490/20-nm EX, 510-nm DCLP, 530/40-nm EM).

Once the neurons were stained by ICC, the same field that had been imaged for FM dye was re-imaged. First, DIC images were acquired for spatial registration of the images for FM and ICC. Next, the signals from VGLUT1 (Alexa Fluor 594), VGAT (Alexa Fluor 488), and MAP2 (Alexa Fluor 405) were acquired, in this order of decreasing excitation wavelengths to minimize photobleaching. They were excited with 595-nm, 490-nm, and 400-nm LED’s (precisExcite; CoolLED), respectively, all at 100% intensity. The corresponding filter cubes had the following properties: 590/55-nm EX, 625-nm DCLP, 665/65-nm EM (“red”); 490/20-nm EX, 510-nm DCLP, 530/40-nm EM (“green”); and 405/40-nm EX, 440-nm DCLP, 470/40-nm EM (“blue”), respectively. Other optical parameters were the same as for FM1-43 imaging.

Eight fluorescence images were acquired using the CCD camera, in a kinetic mode with an exposure time of 1 s each, and the average intensity projection was used for analyses. Image acquisition was controlled by the Solis software (Andor Technology). Images were stored in a SIF format (Andor Technology). The spatial resolution was 0.16 μm/pixel, giving a single full-frame area of 1,392 pixels × 1,040 pixels = 217.7 μm × 162.7 μm. The parameters set for FM and ICC imaging were the same for all imaging fields, to allow for quantitative comparison of staining intensities. The microscope had an internal band-pass filter (340-750 nm) for the fluorescence imaging, and an internal 870-nm LED was used to continuously suppress focus drift resulting from vertical (axial) movement of the microscope stage or imaging chamber (Perfect Focus System; Nikon) by detecting the coverslip-aqueous interface. This anti-focus drift mechanism was activated throughout the experiment, i.e. from the time at which FM dye was imaged until imaging of ICC was completed. Our microscope system enabled us to acquire information from a single imaging field on each coverslip. Extensive care was used to avoid moving the imaging chamber (e.g. mechanically pushing the microscope stage) during any experiment. When apparent chamber displacement was noted, the experimental data were discarded.

### 2.6. Live-cell staining of functional nerve terminals with Ca^2+^ indicator dye and FM dye for full on-stage ICC

Neurons were loaded with a green-emitting [Ca^2+^]_c_ indicator Fluo-5F-AM (F14222; Thermo Fisher Scientific) (Iwabuchi et al., 2013a; Iwabuchi et al., 2013b). Functional nerve terminals were loaded with FM4-64 (T13320; Thermo Fisher Scientific), a red-shifted version of FM1-43, by spontaneous synaptic activities (Iwabuchi et al., 2014a; Kakazu et al., 2012a; Kakazu et al., 2012b). These were achieved by immersing the coverslips in prewarmed Minimum Essential Medium (MEM, 51200-038; Thermo Fisher Scientific) containing 1 μM Fluo-5F-AM and 2.5 μM FM4-64, in the culture incubator at 37°C for 10 min. After staining, the neurons were rinsed by transferring them to dye-free Tyrode’s solution for 10 min at room temperature. The neurons were then transferred to an imaging chamber that was equipped with field-stimulation electrodes (RC-21BRFS; Warner Instruments). The floor of the imaging chamber was a round coverslip (25-mm diameter, thickness 0.09-0.12 mm, No. 0, 633017; Carolina Biological Supply). The neurons were washed with Tyrode’s solution using the bath-perfusion line for 5-10 min. During this washing period, neurons were selected for imaging as described in section 2.5. All subsequent procedures were performed on the microscope stage.

### 2.7. Live-cell fluorescence imaging of neurons for full on-stage ICC

In the partial on-stage ICC (section 2.5), neurons were imaged using a CCD camera (Clara) that is characterized by slow acquisition, low sensitivity but high spatial resolution. In the full on-stage ICC, neurons were additionally imaged using an electron-multiplying CCD (EMCCD) camera (iXon 860; Andor Technology) that is characterized by fast acquisition, high sensitivity but low spatial resolution for analyzing functional dynamics. The EMCCD camera was controlled by the Solis software (Andor Technology), and used in a full-frame format without binning, with a pre-amplifier gain of 4.5, and EM gain of 50 for fluorescence and 0 for DIC, with a 10 MHz pixel readout rate, and without binning. It was cooled at ≤-80°C, with the assistance of a liquid recirculating chiller (Oasis 160; Solid State Cooling Systems, Wappingers Falls, New York, USA). Cells were imaged using the same 40× objective lens (Plan Fluor; Nikon), but with a 0.7× intermediate coupler. The spatial resolution was 0.83 μm/pixel, giving a full-frame area of 128 pixels × 128 pixels = 106.0 μm × 106.0 μm. Both cameras were attached to the same inverted microscope (Eclipse Ti-E; Nikon).

First, DIC images were acquired using CCD and EMCCD cameras. The exposure time for the EMCCD camera was 50 ms, with EM gain of 0. Second, the images of FM4-64 were acquired using CCD and EMCCD cameras. For imaging with the EMCCD camera, the parameters were ten-image acquisitions with an exposure time of 20 ms, EM gain of 50, 595-nm LED intensity of 10% and the “red” filter cube. Third, [Ca^2+^]_c_ signals were acquired using the EMCCD camera. Fluo-5F was excited with 490-nm LED at 10% intensity, and imaged with the exposure time of 7.5 ms, EM gain of 50, the rate of 100 frames per second (i.e. every 10 ms), and the “green” filter cube. Neurons were extracellularly stimulated by an electrical field stimulation of 1-ms duration and 30-mA current intensity, applied through a constant-current, isolated stimulator (DS3; Warner Instruments). Ten background images were acquired under the same condition but with the camera shutter closed. An 8-channel programmable pulse generator (Master-8; A.M.P.I., Jerusalem, Israel) was used to control the Solis software, EMCCD camera, LED, and the electric stimulation. In order to analyze the increase in [Ca^2+^]_c_ signals, an image stack was generated to represent a fold-change [ΔF(t)/F_0_] in Fluo-5F intensity time course [F(t)] from the pre-stimulus baseline level (F_0_), after the averaged background image was subtracted from the raw F(t) images. The [Ca^2+^]_c_ dynamics was measured by assigning regions of interest (ROI’s, 2 × 2 pixels) to neurites in the ΔF(t)/F_0_ image stack, and using the averaged intensity of the pixels as the ROI intensity. A single image for [Ca^2+^]_c_ dynamics was generated by averaging 25 frames in the ΔF(t)/F_0_ image stack near the response peak.

### 2.8. Fixed-cell staining of neurons for full on-stage ICC

After live-cell imaging was completed, the cells were fixed chemically for 15 min, with a total volume of 10 ml. The fixative was the same as in the partial on-stage ICC (section 2.4), but it was applied to the cultured neurons on-stage through the first of the two inlet lines. After fixation, the cells were washed with PBS, for 20-30 min, through the first inlet line and the bath-perfusion line. Subsequent ICC procedures were the same as in partial on-stage ICC (section 2.4), except that both the antibody solutions and their washing solutions were applied through the second inlet line.

### 2.9. Fluorescence imaging of neurons during full on-stage ICC

Imaging conditions with the CCD camera (Clara) were the same as in partial on-stage ICC (section 2.5), including the optics, such as LED’s and filter cubes. Additionally, the EMCCD camera was used for acquiring DIC and fluorescence images. For each fluorophore, ten fluorescence images were acquired with the exposure time of 200 or 300 ms, EM gain of 50, and LED intensity at 100%. The averaged images were used for further analyses.

### 2.10. Correction of lateral translations, for both partial and full on-stage ICC

We used the DIC images of the same structure (e.g. dendritic branches) as a spatial guide for judging the degrees of lateral translation and for correcting them. Standard DIC optics do not require fluorescence filter cube in the optical pathway, i.e. the transmitted light simply travels through the empty “DIC channel” in a filter-cube revolving turret (or slider or anything equivalent with the filter-changing capability). We paired the imaging in one fluorescence channel with the DIC imaging in combination with the same fluorescence filter cube used for the fluorescence imaging. For example, in the partial on-stage ICC, the imaging proceeded in the following order: DIC imaging in “DIC channel” (for standard structural observation) → DIC imaging in green channel → fluorescence imaging in green channel (for FM1-43) → on-stage ICC → DIC imaging in “DIC channel” (for overall alignment after ICC) → DIC imaging in red channel → fluorescence imaging in red channel (for VGLUT1) → DIC imaging in green channel → fluorescence imaging in green channel (for VGAT) → DIC imaging in blue channel → fluorescence imaging in blue channel (for MAP2). Using the first DIC image as a reference for the entire sequence, the later DIC images were translated laterally along the x- and y-axes. The numbers of pixels required to translate the test DIC images and match the reference DIC image were recorded as (Δx, Δy). The same coordinate translations were applied to the paired fluorescence image. The origin of the coordinates is at the top-left corner, with the positive x-axis pointing right and the positive y-axis pointing down.

### 2.11. Image analyses

For overlaying images, the original grayscale images were converted to an 8-bit format, with the minimum and maximum intensity in each field assigned as 0 and 255. The grayscale images were then pseudo-colored and overlaid. For isolating punctate signals from diffusely distributed background noise in fluorescence images, we applied the Difference of Gaussian(s) (DoG) (Iwabuchi et al., 2014b). DoG is based on 1) defining a diffuse background as the image blurred by a two-dimensional Gaussian function, and 2) subtracting it from the original image. The standard deviations (R values in pixels) of the blurring Gaussian functions were 4 for FM4-64, VGLUT1, VGAT, and 5 for MAP2 in CCD images; 2 for FM4-64, Fluo-5F, VGLUT1, VGAT, and 5 for MAP2 in EMCCD images. DoG was not applied to DIC images. All image analyses were performed using the ImageJ software (W. S. Rasband; National Institutes of Health, Bethesda, Maryland, USA) (Schneider et al., 2012).

## 3. Results

### 3.1. Flowcharts of procedures for fixed-cell ICC

Figure 1 shows flowcharts of four general protocols for fixed-cell ICC. They are performed while the specimen is either a) on laboratory benches, i.e. outside the microscope stage (off-stage), or b) on the light-microscope stage, positioned in the optical pathway and ready to be imaged (on-stage) (**Fig. 1**).

The most prevalent ICC protocol (Off-stage ICC #1, **Fig. 1A**) is a conventional one that does not involve live-cell fluorescence imaging. Every procedure is performed outside the microscope stage. Only in the final procedure, the specimen is brought onto a microscope stage for imaging. Key ICC procedures from chemical fixations to treatments with primary and secondary antibodies are indicated by a curly bracket. A standard combined protocol (Off-stage ICC #2, **Fig. 1B**) is for the fixed-cell ICC preceded by live-cell fluorescence imaging. After the live-cell imaging, ICC procedures are performed off-stage. Two sessions of fluorescence imaging (“Image FL”) require spatial alignment of acquired images for integrating information. In one version of our method (On-stage ICC #1 or partial on-stage ICC, **Fig. 1C**), chemical fixation is performed off-stage and subsequent procedures are performed on-stage. Live-cell staining can be done either off- or on-stage, depending on experimental purposes. In another version of our method (On-stage ICC #2 or full on-stage ICC, **Fig. 1D**), all key procedures are performed on-stage. Live-cell staining can be done off- or on-stage. On-stage chemical fixation was reported for cultured neurons, simultaneously in combination with electrophysiological recording (Rosenmund and Stevens, 1997). Our method can be viewed as an extension of the method to ICC.

There are common features among these protocols. First, these ICC protocols are based on indirect immuno-fluorescence method. In this method, an untagged primary antibody is targeted against epitope(s) of an antigen. A secondary antibody is targeted against the primary antibody, and is conjugated with a fluorescent probe (fluorophore). Second, our antigens are located intracellularly. This condition requires that the cells become fixed. For this purpose, we used chemical fixation by aldehyde-based fixatives. Third, the intracellular location of antigens additionally requires us to apply detergents, such that the plasma membrane, and organellar membrane additionally (depending on the antigen localization), are permeabilized for easy access of antibodies to antigens.

The two on-stage ICC protocols include live-cell staining with fluorescent probes for neuronal functions. A difference of these on-stage protocols from the off-stage protocol is that they are interspersed with imaging sessions. In these sessions, the specimens are imaged for the fluorescent signals that originated from fluorophores representing live-cell functions or from fluorophores conjugated to secondary antibodies indicating fixed-cell ICC signals. Critically, the specimens are imaged using transmitted light, e.g. DIC optics, in combination with the fluorescence filter cubes used for fluorescence imaging. The transmitted-light imaging mode can record spatial displacements that could have happened during any part of the previous steps, including pushing a microscope stage by mistake, and an improper alignment of fluorescence filter cubes in the optical path.

### 3.2. Image registration using DIC optics

We will demonstrate representative results based on the full on-stage ICC protocol (**Fig. 1D**) (N = 3 cultures). We used two cameras to maximize the efficiency of acquiring different types of information. One camera (CCD, Clara) was suitable for high spatial resolution for morphological information. The other camera (EMCCD, iXon) was suitable for fast acquisition of functional [Ca^2+^]_c_ signals but with low spatial resolution. Cultured mouse hippocampal neurons were stained off-stage with a Ca^2+^ indicator Fluo-5F-AM for monitoring [Ca^2+^]_c_, and with FM4-64 to label functional nerve terminals. Stimulus-induced dynamics of [Ca^2+^]_c_ was imaged and the presynaptically silent synapses were independently evaluated.

DIC images were taken at four steps of the full on-stage ICC protocol, indicated by “Image FL & DIC (with FL filters)” in **Fig. 1**. They are shown as the pseudo-colored overlay of a test image at a certain step (green) and of a reference image under the live-cell condition (red) (**Fig. 2**). The test images were acquired, immediately after completing chemical fixation (**Fig. 2A**), immediately after membrane permeabilization with Triton X-100 (**Fig. 2B**), and after completion of ICC (**Fig. 2C**). Displacements of the test image from the reference image are visible as red or green regions. After lateral translations, i.e. moving the test image along the x- and y-axes by (Δx, Δy) pixels, the test image matched the reference image, with no displacements indicated by yellow color.

**Fig. 2.**
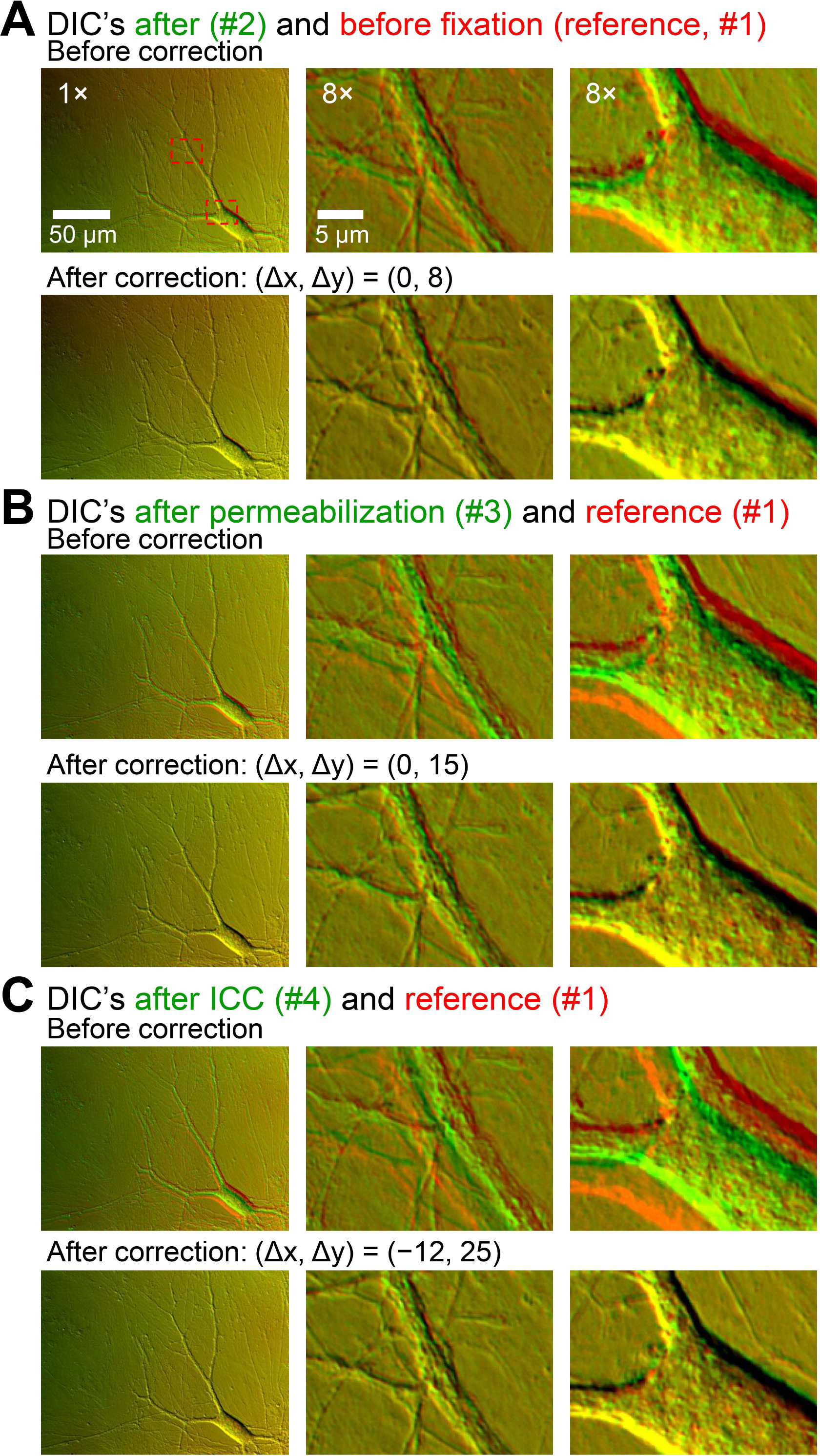
DIC-based corrections of image displacements during the full on-stage ICC. Images were acquired using a CCD camera with high spatial resolution but slow acquisition. Cultured mouse hippocampal neurons were imaged with DIC optics. All panels represent overlays of two images. The DIC image acquired during live-cell imaging (image #1) was used as a spatial reference image and was pseudo-colored red in all panels. DIC images were further acquired during later procedures: immediately after completing chemical fixation (image #2, **A**), immediately after membrane permeabilization (image #3, **B**), and after completion of ICC procedures (image #4, **C**). These test images were pseudo-colored green. In **A, B, C**, top and bottom rows represent overlays before and after correcting the displacements, respectively. Displacements between two images appear as red or green, and no displacements appear yellow. The degree of displacement, and therefore the degree of required correction, is indicated by how much the test image had to be translated laterally along the x- and y-axes to match the reference image [(Δx, Δy) in pixels]. For a simple demonstration, the illustrated images were selected from the ones acquired without fluorescent filters.

These results show that only a small degree of corrections was necessary to match the test image to the reference image (up to several tens of pixels in most of the experiments). They also show that the corrections were limited to lateral translations, not involving any image rotations. In contrast, when the off-stage ICC protocol (**Fig. 1B**) was used, it was very rare for us to have the test image covering the same field as the reference image. Even when the target cells were in the same field, corrections required a large degree of both lateral translations and rotations (data not shown).

The neuronal cultures included soma and well-developed neurites (dendrites and axons). Somata tended to become distorted by the aldehyde-based chemical fixation, which was visible as changes in somatic sizes and shapes. In some imaged fields, parts of neurites became distorted, with disproportionate displacements and with changes in curvature. When these non-linear changes were noticed in the neurites, the data were discarded, because our experiments were focused on functions and structures of dendrites where synapses formed.

### 3.3. Fluorescence image registration

After image displacements were corrected using DIC images, the same translation parameters (Δx, Δy) were applied to fluorescence images obtained at the same steps. These transformations corrected the displacements in fluorescence images. We overlaid the following fluorescence images onto the live-cell, reference DIC image: purely morphological information (VGLUT1, VGAT, MAP2, **Fig. 3A**), functional and morphological information of glutamatergic nerve terminals (FM4-64, VGLUT1, MAP2, **Fig. 3B**), and the corresponding information of GABAergic nerve terminals (FM4-64, VGAT, MAP2, **Fig. 3C**). Yellow and red arrows in **Figs. 3B** and **3C** point at some of the active (i.e. positive for FM4-64) and silent nerve terminals (i.e. negative for FM4-64), respectively. Accurate alignments of the fluorescence images demonstrated not only the functionally-labeled, active nerve terminals, but also the unlabeled, presynaptically silent nerve terminals.

**Fig. 3.**
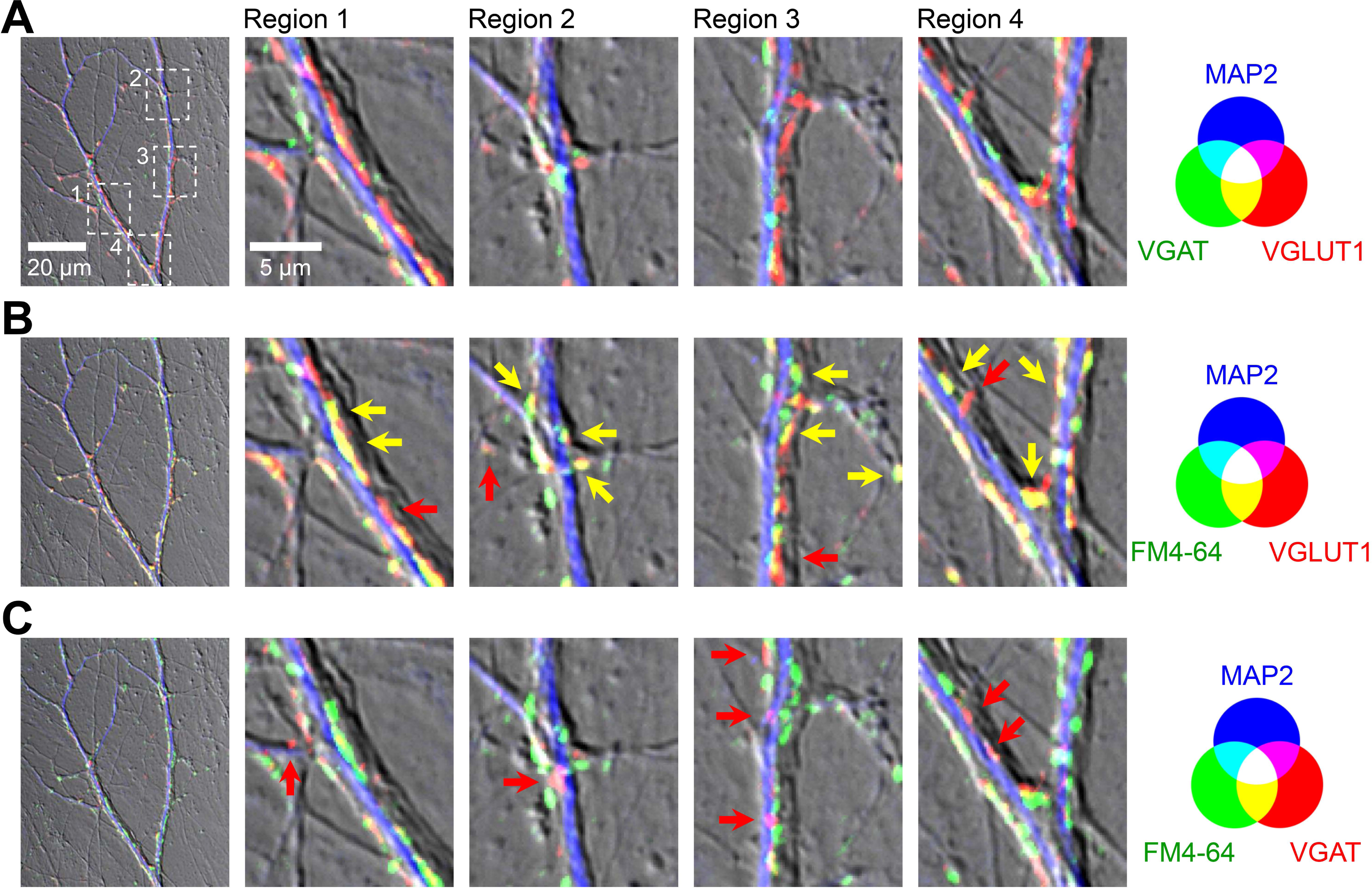
Overlay of functional and morphological signals acquired by the full on-stage ICC. Images were acquired using a CCD camera with high spatial resolution but slow acquisition. Functional signal was the staining intensity of FM4-64 dye which was taken up by recycling synaptic vesicles in functionally active nerve terminals. Morphological signals were derived from ICC for markers of: glutamatergic nerve terminal, vesicular glutamate transporter 1 (VGLUT1); GABAergic nerve terminal, vesicular GABA transporter (VGAT); and dendrites, microtubule-associated protein 2 (MAP2). Different combinations of the four fluorescent signals were overlaid on the reference DIC image. (**A**) VGAT (green), VGLUT1 (red) and MAP2 (blue). (**B**) FM4-64 (green), VGLUT1 (red) and MAP2 (blue). (**C**) FM4-64 (green), VGAT (red) and MAP2 (blue). In **B** and **C**, some of the presynaptically active and silent synapses were labeled with yellow and red arrows, respectively. The imaged field corresponds to a part of the upper half of the field in **Fig. 2**.

### 3.4. Image registrations using a fast-acquisition EMCCD camera

**Figures 2** and **3** were acquired using a CCD camera with high spatial resolution. In order to analyze functional [Ca^2+^]_c_ dynamics, additional images were acquired using an EMCCD camera with fast acquisition. Displacements of DIC images were corrected, as in **Fig. 2** (**Fig. 4**). Note that the number of pixels required for corrections, (Δx, Δy), was much less (After correction, **Fig. 4B**), because the image resolution of the EMCCD camera was much lower than that of the CCD camera.

**Fig. 4.**
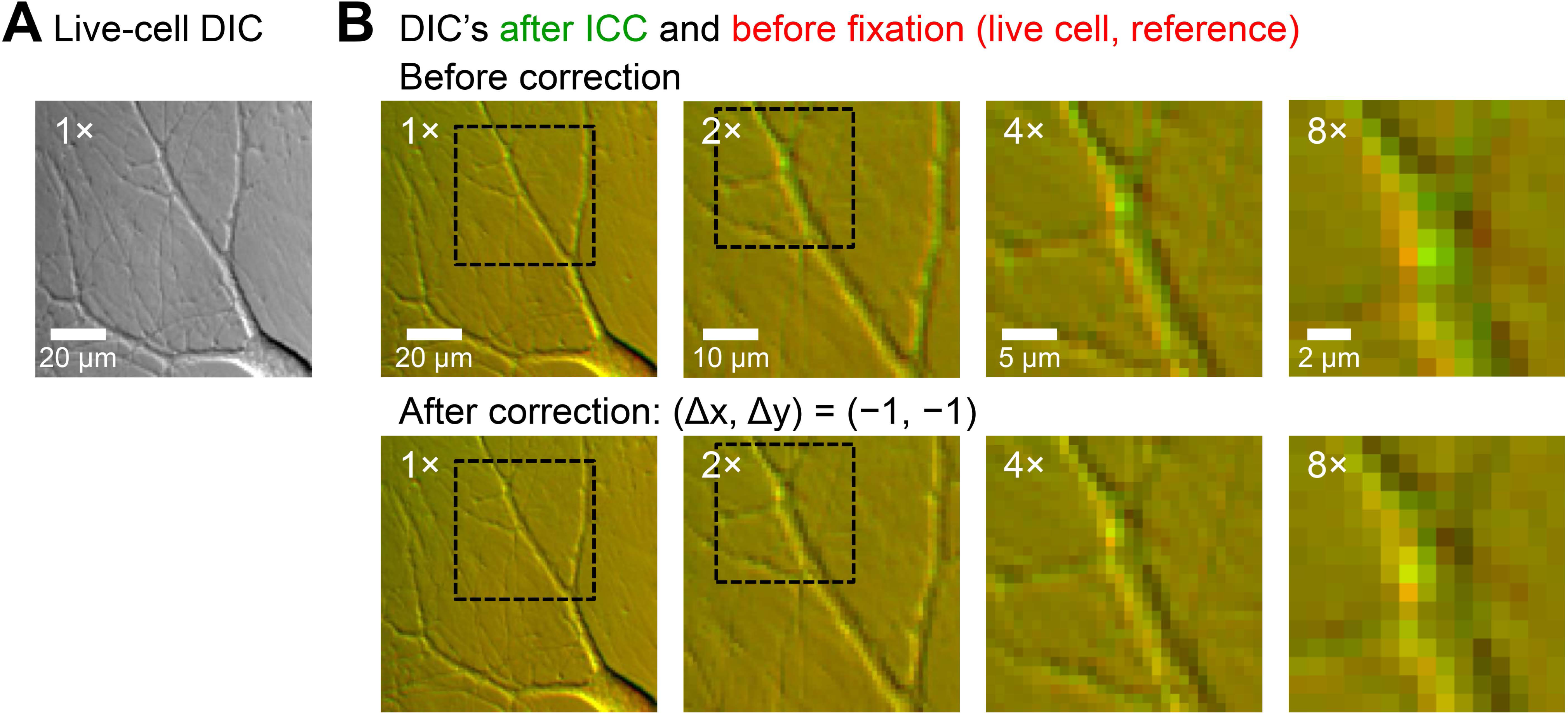
DIC-based corrections of image displacements during the full on-stage ICC. Images were acquired using an EMCCD camera with fast acquisition but low spatial resolution. (**A**) DIC image acquired before chemical fixation. (**B**) Overlays of the reference DIC image acquired during live-cell imaging (red) and the test DIC image acquired after completion of ICC procedures (green). The latter image was not corrected for displacement (top row), and was corrected (bottom row). For a simple demonstration, the illustrated images were selected from the ones acquired without fluorescent filters.

Using the same translation parameters, the fluorescence images were corrected (**Fig. 5**). In the same image field, we could locate all the signals of DIC (**Fig. 5A**), FM4-64 (**Fig. 5B**), locations of peak [Ca^2+^]_c_ responses (**Fig. 5C**), locations and dynamics of [Ca^2+^]_c_ responses (**Fig. 5D**)VGLUT1 (**Fig. 5E**), VGAT (**Fig. 5F**) and MAP2 (**Fig. 5G**).

**Fig. 5.**
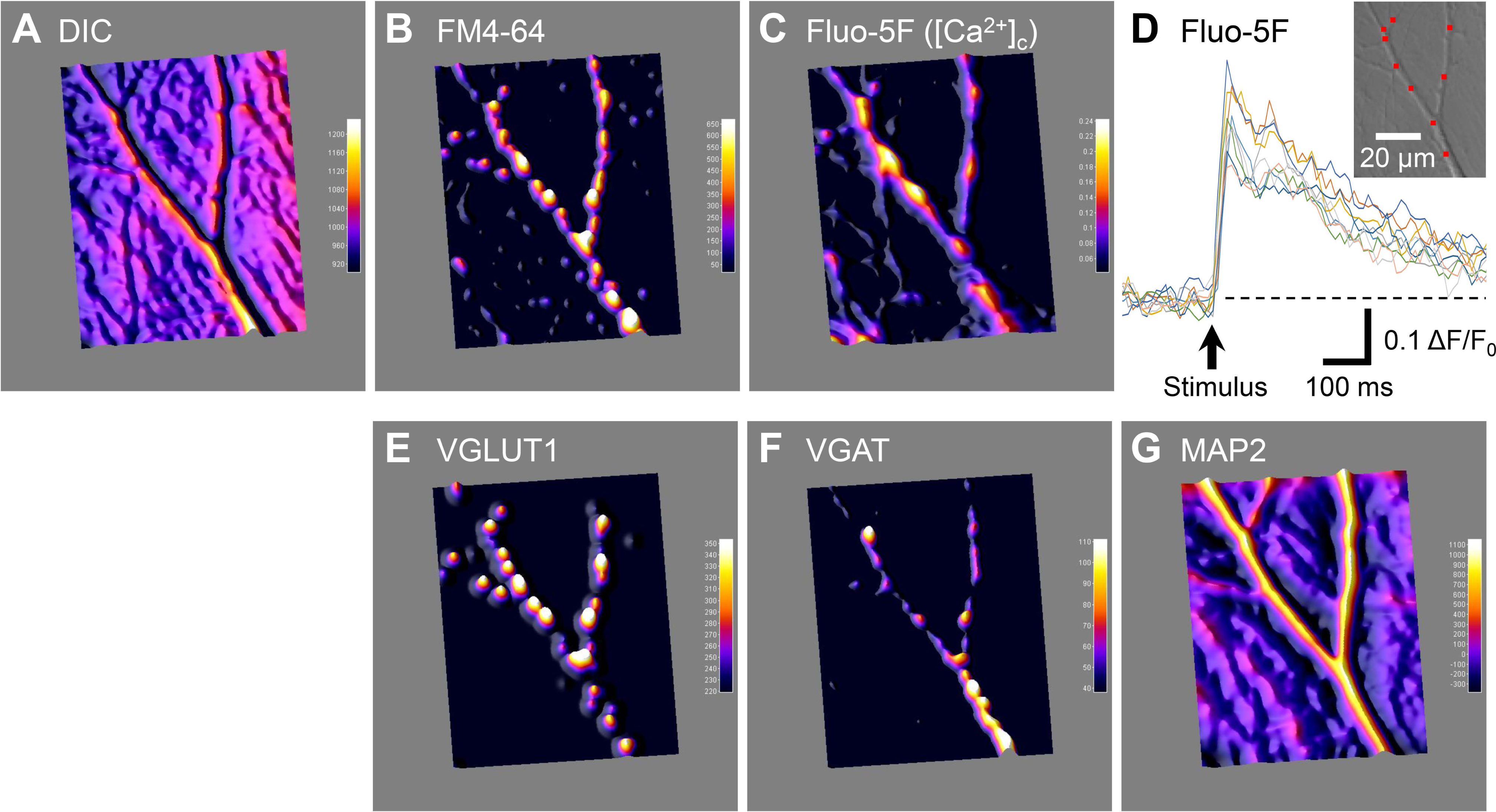
Functional and morphological signals after image displacements were corrected. Images were acquired using an EMCCD camera with fast acquisition but low spatial resolution. Functional signals were the staining intensity of FM4-64, and [Ca^2+^]_c_ dynamics. Panels **A-C** and **E-G** are surface plots: (**A**) reference DIC image acquired during live-cell imaging, (**B**) FM4-64, (**C**) Fluo-5F, (**E**) VGLUT1, (**F**) VGAT, (**G**) MAP2. (**D**) [Ca^2+^]_c_ dynamics. Nine overlaid traces represent a fold-change in Fluo-5F intensity [ΔF(t)/F_0_], in response to a single electrical stimulation (arrow). Individual traces were obtained from nine regions of interest (ROI’s, 2 × 2 pixels) shown in the inset. The image field was the same as in **Figs. 2-4**.

We independently performed experiments with the partial on-stage ICC protocol. Live cultured hippocampal neurons were stained with the FM dye, and were fixed and stained by ICC for VGLUT1, VGAT and MAP2 (i.e. without [Ca^2+^]_c_ imaging) (N = 3 cultures). These results of presynaptically active and silent synapses were similar to the results presented above with the full on-stage ICC protocol, and thus the data are not shown.

### 3.5. Spatial resolution of light microscope

Two questions are relevant to the spatial resolution in our imaging experiments. First, how small / large is the resolution with respect to the size of biological structures? Second, to what extent is the resolution constant across the image field? To answer these questions, we evaluated the spatial resolution under a condition similar to that for functional and morphological imaging using EMCCD camera (fast acquisition but of low resolution) (**Figs. S1, S2**). We first measured the point-spread function (PSF), i.e. the spatial distribution of fluorescence emitted from a sub-resolution, point source. Then the resolution was defined as a width of the PSF at 50% peak height (full width at half maximum, FWHM) (Egner and Hell, 2006). Similar PSF shapes indicated that the spatial resolution was constant across the image field. This notion was supported by the very small variations in the measured resolution: the averaged FWHM was 0.618 ± 0.008 μm (mean ± sem, n = 5 beads). FWHM value was smaller than the size of nerve terminals (∼1 μm). The pixel size of EMCCD camera was ∼0.83 μm, indicating that a pixel interpolation associated with image rotations will cause a large effect.

## 4. Discussion

### 4.1. Summary

We have established a method for spatially correlating fluorescent signals acquired from live-cell functions with those acquired from fixed-cell morphologies, using a widefield optical microscope. This was effective at the micrometer scale, i.e. at the subcellular level of e.g. dendrites and nerve terminals. The method is based on two simple procedures: 1) performing a subset or all ICC procedures on-stage, which confines image displacement to a small degree of lateral translations and no rotations, and 2) correcting residual displacements in fluorescent images, using, as landmarks, the subcellular structures imaged with transmitted-light optics in combination with fluorescence filter sets. The DIC optics played an important role. It was used at least for five versatile purposes: 1) evaluate the health condition of the live neuronal culture, 2) select a culture field to be imaged for experiments, 3) examine whether the imaged field was distorted by chemical fixation, 4) serve as a reference image and provide test images for correcting image displacements, 5) provide a structural image for overlaying fluorescence images that represent neuronal functions and morphology. The effectiveness of these procedures was demonstrated using cultured neurons as an example.

### 4.2. Correlative microscopy

Insight into the biology of cells can be obtained by imaging the same specimen using multiple microscopy modalities and integrating the information from them. Such correlative microscopy sheds light on spatial relationship and correlations of various signals.

In a narrow sense, correlative microscopy refers to correlative (also referred to as correlated) light and electron microscopy (CLEM) (de Boer et al., 2015), a method that integrates imaging information from light and electron microscopy of the same specimen (Ganeva and Kukulski, 2020; Harata et al., 2001). One of the early reports of such an approach appeared (Bopp-Hassenkamp, 1959) soon after electron microscopy was first applied to biological specimens.

In a broad sense, correlative microscopy refers to any combination of microscopy techniques (Loussert Fonta and Humbel, 2015), such as super-resolution fluorescence microscopy [e.g. stimulated emission depletion (STED) microscopy and photoactivated localization microscopy s(PALM)] (Bernhardt et al., 2018; Watanabe et al., 2011), fluorescence imaging *in vivo* (intravital microscopy) (Karreman et al., 2016), microscopic X-ray computed tomography of excised tissue (Karreman et al., 2017), transmission electron microscopy (TEM) (Fuest et al., 2018), atmospheric or environmental scanning electron microscopy (SEM) (Peckys and de Jonge, 2015; Sato et al., 2019), cathodoluminescence of nanoparticles in SEM (Glenn et al., 2012), cryo-electron tomography (Tao et al., 2018), array tomography of electron microscopy (Burel et al., 2018; Collman et al., 2015; Micheva and Smith, 2007), X-ray holography, X-ray scanning diffraction (Bernhardt et al., 2018), X-ray fluorescence imaging and mass spectrometry imaging (Decelle et al., 2020). Some authors emphasized this integration of information from multiple imaging methods, by proposing a “morphomics” notation (Lucocq et al., 2015). Correlative microscopy should also include light-microscopy information of the same specimen, as a bridge between live- and fixed-cell conditions. Thus we view our protocol as one small component of correlative microscopy in a broad sense.

### 4.3 Methods for spatial registration in correlative microscopy

Overall, there are at least four categories of approaches to enable alignment of multiple images in correlative microscopy.

The first approach is based on the creation of fiducial markers, i.e. landmarks that have both optical and electron contrast. It can involve: etching of glass coverslips with a diamond knife (Darcy et al., 2006b); etching of glass coverslips with a specific grid pattern (e.g. CELLocate); printing of coverslips with gold nanoparticles (Benedetti et al., 2014; Prabhakar et al., 2018) or titanium (Rodighiero et al., 2015). It can also involve applying the following onto specimens: single- or multi-color fluorescent beads (Schorb et al., 2017; Watanabe et al., 2014; Watanabe et al., 2011); gold particles (Watanabe et al., 2014); fluorescently labelled gold particles (Mohammadian et al., 2019); quantum dots and Y_2_O_3_:Eu^3+^ nanoparticles (van Hest et al., 2019). Furthermore the approach includes introducing artificial autofluorescence to surrounding tissue with or without photo-oxidation (near-infrared branding) (Bishop et al., 2011; Karreman et al., 2017; Lees et al., 2017; Takahashi-Nakazato et al., 2019). The fiducial-marker approach can address procedural Problems 1 and 2 described in the Introduction, provided that the fiducial markers are applied at a density high enough to ensure that they appear in any imaging field.

The second approach uses a fluorophore or molecular probe that is visible by both fluorescence imaging and another imaging modality, such as TEM and SEM. Examples include quantum dots (Hanifi et al., 2019), which can be conjugated to antibodies (Giepmans et al., 2005; Killingsworth and Bobryshev, 2016; Peckys and de Jonge, 2015), and a fluorophore and a gold cluster conjugated to antibodies (FluoroNanogold) (Takizawa et al., 2015). This approach has multiple variants. In one, the secondary antibody solution in ICC is a mixture of antibodies that are conjugated with fluorophores and with gold particles (Rodighiero et al., 2015). In a second variant, non-fluorescent gold nanoparticles are visualized by local surface plasmon resonance of light microscopy and also by electron microscopy (Haruta et al., 2019). In a third variant, an electron-dense product is generated from fluorophores through a process termed photo-oxidation or photoconversion (Bishop et al., 2011; Darcy et al., 2006a; Dobson et al., 2019; Harata et al., 2001; Hoopmann et al., 2012; Opazo and Rizzoli, 2010; Schikorski, 2010, 2014). The use of a special fluorophore is not always necessary for live and fixed cells, with the need depending on the study design. For example, when presynaptically active synapses (as opposed to silent ones) are a research target, they can be identified based on both function (using an FM dye) and structure (using ICC for a nerve terminal marker) (Micheva et al., 2006). In such cases, colocalization of fluorescent signals can be used a method for spatial registration between different fluorescent channels.

The third approach aims at minimizing the need for image realignment by integrating specimen-preparing equipment and/or imaging modalities within a single instrument. Examples include the microtome-integrated light microscope for TEM (Lemercier et al., 2017), microtome-integrated confocal and scanning probe (e.g. atomic force) microscope (Mochalov et al., 2017), cryo-fluorescence and cryo-TEM (Schorb et al., 2017), cryo-fluorescence and cryo-SEM tomography (Gorelick et al., 2019), and fluorescence imaging and environmental SEM of liquid-state cells (Peckys and de Jonge, 2015; Sato et al., 2019). Some of these integrated microscopes are commercially available and might achieve the goal of minimizing or eliminating the need for spatial registration. However, they are resource-consuming (expensive).

The fourth approach is based on software that corrects the displacements of images once they have occurred. This approach is not necessarily independent of the previous approaches. In fact, it can be integrated with any of them. Notably, some software packages can perform spatial registration of three-dimensional data sets (e.g. Amira Software, Thermo Fisher Scientific Inc.) (Karreman et al., 2017), and of distinct images based on analogies (Cao et al., 2014).

### 4.4. Advantages of our system

Our system works efficiently in experiments where the necessary spatial resolution is at the subcellular, micrometer level. The advantages are that this system is applicable even when the live- and fixed-cell fluorescence signals do not colocalize spatially (as in presynaptically silent synapses), and when the live-cell fluorescence signals are transient and local, and thus invisible by fixed-cell imaging (as in [Ca^2+^]_c_ imaging). Furthermore, it does not require a change of fluorophores, the application of fiducial markers, or modification of existing microscopes or experimental systems in the laboratory, provided that the basic steps of the experiments can be performed individually. Also, our method requires neither a deep budget nor deep knowledge of optics. Note that our method is not designed for accuracy at the molecular, nanometer scale which requires super-resolution fluorescence microscopy or electron microscopy.

### 4.5. Points of improvement

Our approach is based on manual manipulations, which can be labor-intensive. Thus it is not feasible to analyze a large number of data sets. It can be improved using some of the technical features that are available for other forms of correlative microscopy as a guide. During image acquisitions, the efficiency and accuracy of spatial registration will be improved if it incorporates a computer-controlled motorized microscope stage, in combination with accurate measurement of the x and y coordinates of high-density fiducial markers. After images are acquired, residual displacements will be corrected using image-alignment software in an objective manner. An example of image translation software, available as freeware, is the ImageJ plugin MultiStackReg (http://bradbusse.net/sciencedownloads.html; accessed on 6/30/2020). It allows for two or more images to be aligned using one channel, saves the parameters of image transformations, and then uses them to align the images in the other channels, provided the lateral displacement and re-orientation are not extensive. It might also be possible to realign images based on analogies among images acquired by different imaging modalities (Cao et al., 2014). Overall, further improvements are expected to lead to semi-automated image analyses.

From the biological perspective, two features have not been addressed in this work. One feature is a slight morphological change caused by aldehyde-based chemical fixation, i.e. tissue shrinkage and morphology distortion. It is unavoidable under the current protocol, and affects some cellular structures and imaged fields more markedly than others. For example, whereas somata showed a large degree of change, some neurites were spared. There is a possibility that the shrinkage and distortion are partly corrected using imaging software, but such non-linear transformation could itself introduce unexpected artifacts. Thus in the current work, we focused on image fields that were stationary after lateral translations, and discarded data if distortion was apparent. The system could be improved by incorporation of a fixation method that maintains both cellular morphology and antigenicity in ICC procedures.

Another feature that has not been addressed is a choice of specimens. Cultured cells were used as an exemplar specimen in this study. Our on-stage ICC protocols will be effective, when the structural landmarks are easily visible with the transmitted-light optics and used as a spatial reference. Thus a monolayer culture, such as our primary neuronal culture with extensive neurites, is an optimal specimen. However, other specimens can be used if natural landmarks are available. For example, blood vessels in brain slice preparations might serve as structural landmarks, when the plasma membrane of endothelial cells is labeled with a lipophilic carbocyanine dye DiI (Konno et al., 2017; Li et al., 2008).

With modifications and improvements, the method described here will be applicable to a wide variety of light-microscopy experiments involving subcellular structures and organelles, in neuroscience and other biomedical fields.

## ABBREVIATIONS

[Ca^2+^]_c_: cytosolic Ca^2+^ concentration
CCD: charge-coupled device
CLEM: correlative light and electron microscopy
DIC: differential interference contrast
DoG: difference of Gaussian(s)
EMCCD: electron-multiplying CCD
F(t): time course of fluorescence intensity
F_0_: pre-stimulus fluorescence baseline level
ΔF(t)/F_0_: time course of fold-change in fluorescence intensity
ICC: immunocytochemistry
LED: light-emitting diode
MAP2: microtubule-associated protein 2
NGS: normal goal serum
PALM: photoactivated localization microscopy
PBS: phosphate-buffered saline
PSF: point-spread function
SEM: scanning electron microscopy
STED: stimulated emission depletion
TEM: transmission electron microscopy
VGAT: vesicular GABA transporter
VGLUT1: vesicular glutamate transporter 1
(Δx, Δy): coordinate translation required to correct an image displacement.

## Acknowledgements

The authors thank Dr. Kristina Micheva (Stanford University) for comments about image alignment software. This work was supported by the US Department of Defense (W81XWH-14-1-0301, W81XWH-18-1-0136 to NCH) and Dystonia Medical Research Foundation (to NCH).

## SUPPLEMENTARY INFORMATION

## Supplementary Materials and Methods

*Comments on the experimental procedures*

- Selection of FM dyes and Ca^2+^ indicator dyes
  - Representative dyes for labeling the recycling synaptic vesicles include green-emitting FM1-43 and red-emitting FM4-64 (Iwabuchi et al., 2014). In the partial on-stage ICC protocol, either FM dye can be used interchangeably, by using proper light sources and fluorescent filters. In fact, different FM dyes can be used to differentially label different synaptic vesicle pools in a single experiment (Klingauf et al., 1998; Richards et al., 2000).
  - In the full on-stage ICC protocol, it is possible to use a different pair of FM dye and Ca^2+^ indicator dye from the one described above, as long as the fluorescence emission from one dye exhibits minimal bleedthrough into the detection channel of another dye. However, emission spectra overlap is considerably smaller with a pair of green-emitting Fluo-5F and red-emitting FM4-64 than with a pair of green-emitting FM1-43 and red-emitting Ca^2+^ dye, such as Asante Calcium Red (Hyrc et al., 2013). This is because FM dyes exhibit broad emission spectra in comparison to Ca^2+^ dyes (vendor information, Spectra Viewer; Thermo Fisher Scientific).

- Negligible effects of FM dyes and Ca^2+^ indicators on ICC results
  - The emission spectra of FM1-43 and FM4-64 were wide, and overlapped extensively with those of the Alexa fluorophores conjugated with secondary antibodies (Spectra Viewer; Thermo Fisher Scientific). However, this spectral overlap did not affect our experiments, because FM dyes were lost from the specimen during permeabilization with 0.1% Triton X-100 (Kawano et al., 2012; Moulder et al., 2010), and therefore the FM signals were absent by the time when ICC results were imaged.
  - Ca^2+^ indicators can be lost similarly during permeabilization. Even if a small amount of the dyes remains, the residual signal from the dyes was spatially diffuse when the neurons were chemically fixed at the resting state of [Ca^2+^]_c_ (F_0_). Diffuse signals are effectively eliminated by DoG for the fluorescence signal of ICC (VGAT in the green channel in the demonstrated experiment). Note that the regional selectivity of [Ca^2+^]_c_ responses (**Fig. 5C**) reflects a small response with the peak value of ΔF/F_0_ ∼30%.

- General comments about ICC procedures
  - Glutaraldehyde was used at 0.2% for our cultured neurons, as reported previously (Moulder et al., 2010). Its use at a high concentration of 1.25% (Clancy and Cauller, 1998) or 2% (Harata et al., 2001) would be accompanied by significant autofluorescence, which needs to be attenuated, e.g. by sodium borohydride (NaBH_4_) (Clancy and Cauller, 1998; Tagliaferro et al., 1997), and by ammonia-ethanol or Sudan Black B (Baschong et al., 2001). But at the concentration range of 0.2-0.8% used in the current study, this was not an issue. Use of a high concentration of glutaraldehyde is also a concern because it can reduce the antigenicity of some antigens.
  - Blocking step by normal goat serum (NGS) was combined with the permeabilization step by Triton X-100, by dissolving Triton X-100 in the blocking solution (PBS with NGS). They can be separated into two steps, with permeabilization preceding the blocking.
  - The antibodies were dissolved in the blocking solution. They can additionally have Triton X-100.
  - Spectral bleedthrough between pairs of fluorophores conjugated to the secondary antibodies was negligible. The intensity was as weak as the background fluorescence intensity for a negative control in which primary antibodies were omitted from ICC (data not shown). Thus the acquisition of one signal did not interfere with that of another.

- Manipulations that contributed to increasing the ICC signal intensity
  - Treatment with primary antibodies was for 3 hours, because this provided stronger signal than treatment for shorter periods, e.g. 2 hours. The dilution factor of primary antibodies was 1:2000, but 1:1000 yielded similar results, with a slightly stronger signal. Naturally, the dilution factors should be determined empirically for each antibody that the experimenters use in their protocols.
  - During treatment with primary antibodies, it was important to introduce some turbulence to the solution in the chamber bath. This was required for good mixing of the solution in the imaging chamber and to ensure that the antibodies had access to the antigens. In our system, ICC solutions were applied through the inlet lines manually, using syringes dedicated to individual solutions/lines. For each turbulence, 300-500 μl of the primary antibody solution were pulled from the imaging chamber through the inlet line using a syringe, and approximately the same volume was gently pushed back into the chamber. This pull-push turbulence took ∼5 sec and was repeated for ∼1 min, every ∼30 min during the treatment period. Without this turbulence, the fluorescence signals were weak.
  - Treatment with secondary antibodies was for 60-90 min. This provided stronger signals than treatment for shorter time periods, e.g. 45 min, although the effect was not large.
  - During treatment with secondary antibodies, the pull-push turbulence was introduced once or twice during the treatment period. This manipulation was not as important as during the treatment with primary antibodies.
  - Alexa dyes were used as the fluorophores conjugated with secondary antibodies. Under the current experimental conditions, they provided much brighter signals than quantum dots, another set of common fluorophores that we tried, e.g. F(ab’)2-goat anti-rabbit IgG secondary antibody conjugated with quantum dot (Qdot) 525 (Q11441MP; Thermo Fisher Scientific) or F(ab’)2-goat anti-mouse IgG secondary antibody conjugated with Qdot 605 (Q11002MP; Thermo Fisher Scientific). The weak signals of quantum dots were most likely due to their large size and the associated difficulty of entering into cells for accessing intracellular antigens under the current experimental conditions.

- Solution flow
  - Except for the bath perfusion, all solutions were applied through the two inlet lines manually, using syringes dedicated to individual solutions.
  - Extreme care was taken to avoid directing solution outflow from the two inlet lines toward the cells while ensuring that solution flowed smoothly through the bath. Otherwise, the cells were easily disturbed or blown away.
  - Care was taken to avoid the tips of the inlet lines from touching the bottom of imaging chamber, i.e. to leave some vertical space. Otherwise, the pull-push turbulence could lead to drying up the cells.
  - The pull-push turbulence was also applied during chemical fixation of full on-stage ICC.
  - It is important to carefully control the suction system for avoiding cells being dried up and avoiding overflowing the chamber. It is also important for maintaining efficient solution exchanges, such that the concentration of key solutions (e.g. antibodies) is not affected by dilutions. This is because the volumes of the chamber and of the key solutions (e.g. fixative, antibodies) are limited. For example, when the primary antibodies are applied to the cells, we will proceed as follows: keep the suction line open, stop the bath perfusion for PBS, stop the infusion of blocking solution through the first inlet, start infusion of the antibody through the inlet while the suction system open (because there is a dead volume in the lumen that still contains the previous solution, the blocking solution in this case), close the suction when the total volume of antibody solution has been pushed out using a syringe, and leave the condition at the level during the expected treatment time.
  - Summary of the solution lines in the partial on-stage ICC protocol. The bath perfusion line had only PBS throughout the experiment. One (1^st^) inlet line was for: 1) Triton X-100 in the blocking solution, 2) primary antibodies in the blocking solution, 3) blocking solution (wash-out of Triton X-100 and of primary antibodies). Another (2^nd^) inlet line was for: 1) secondary antibodies in the blocking solution, 2) PBS without NGS (wash-out of secondary antibodies).
  - Summary of the solution lines in the full on-stage ICC protocol. The bath perfusion line was for: 1) Tyrode’s solution (for live-cell [Ca^2+^]_c_ imaging), 2) PBS (for fixed-cell ICC), switched by a three-way stop cock. One (1^st^) inlet was for: 1) fixative, 2) PBS (wash-out of fixative). Another (2^nd^) inlet line was for: 1) Triton X in the blocking solution, 2) primary antibodies in the blocking solution, 3) blocking solution (wash-out of Triton X and of primary antibodies), 4) secondary antibodies in the blocking solution, 5) PBS without NGS (wash-out of secondary antibodies). In other words, these solutions were the same as those applied through two inlet lines in the partial on-stage ICC protocol.
  - Only two inlet lines were used due to the limited space near the imaging chamber. If space is allowed, it would be better to use three inlet lines for the full on-stage ICC protocol. This will separate the primary and secondary antibody solutions into two lines as in the partial on-stage ICC protocol, still isolating one line dedicated to the fixative and its washout.

- The ICC protocol was modified from the one reported earlier, in the work of presynaptically silent synapses (Moulder et al., 2010). The major modifications were as follows:
  - We omitted the use of an FM-dye remover ADVASEP-7, a β-cyclodextran derivative (Kay et al., 1999), during FM dye washout. This was because it can lead to the loss of FM from positively stained structures (Moulder et al., 2010) and it can have deleterious effects on cellular morphology.
  - ICC was performed on-stage.
  - In the original protocol, FM dye was imaged after ICC was completed (Moulder et al., 2010). In order to retain the FM dye in the positively stained structures as much as possible, cellular membranes were permeabilized with Triton X-100 at a low concentration (0.04%) (Moulder et al., 2010). In contrast, in the current protocol, the on-stage ICC allowed us to image the FM dye before permeabilization, i.e. without any FM loss, and also to permeabilize the membrane with Triton X-100 at a high concentration (0.1%) for a stronger ICC signal and without FM signal contamination.
  - ICC was used to detect three antigens per specimen (instead of one or two). This required an additional space on fluorescence spectra, and was made feasible by the prior and complete loss of FM dye from the specimen.
  - Cellular imaging in a mounting medium (Fluoromount-G) was replaced by imaging in aqueous solution (PBS) immediately after ICC staining, because such an aqueous solution provides a refractive index similar to the Tyrode’s solution used during live-cell imaging.

- Washing the experimental system after the imaging experiments are completed
  - It is important to wash the system with care. All tubings and syringes should be cleared of their contents and rinsed with distilled water, e.g. 50-100 ml or more for the bath-perfusion line and 15-20 ml or more through each of the two inlet lines. After rinsing with distilled water, dry up the lumens by pushing through air. Imaging chamber needs to be rinsed extensively as well, e.g. with the running tap water and then with distilled water. Fixative solution remaining in an imaging chamber will affect the quality of the subsequent live-cell experiments. It might be helpful to devote one chamber to live- and fixed-cell experiments, and separate it from chambers individually devoted to experiments for live cells alone and for fixed cells alone.
  - If different combinations of antibodies and fluorophores are applied frequently in different experiments, it might be worth incorporating, into the washing procedures, additional processes for actively removing residual antibodies. Such processes include rinsing with detergents, antibody elution (Micheva et al., 2010) or antibody stripping (Carroll et al., 1999), although we have not tried them.

- Lateral translations of images
  - “Image FL & DIC” step (**Fig. 1C, D**) can be essential or non-essential (and therefore eliminated), depending on the goals of experiments. The reference image and the test images after ICC are essential.
  - Reference image can be chosen arbitrarily based on the experimental goal. In the demonstrated data for the full on-stage ICC protocol, the reference image was acquired before chemical fixation. But in the partial on-stage ICC protocol, the reference image was acquired immediately after chemical fixation and before permeabilization.
  - It is not absolutely necessary in these protocols to perform the standard DIC imaging, i.e. plain DIC without fluorescence filters. However, such DIC imaging is generally recommended for recording structural information of the imaged field.
  - In the section 2.10, we described the most comprehensive sequences of DIC-fluorescence pairs. If the degrees of image shifts associated with filter cube rotations remain constant among experiments, it might be possible to simplify the procedures by removing the DIC imaging with filter cubes, but still retaining the DIC imaging without any filter cube.

### Visualization of point-spread function (PSF) analyzed with sub-resolution fluorescent beads (for Supplementary Figures)

Optical resolution was evaluated by measuring the spatial distribution of fluorescence emitted from a sub-resolution, point source (PSF). The solution suspending green fluorescent beads of the PS-Speck Microscope Point Source Kit (175-nm diameter, P7220; Thermo Fisher Scientific) was dried on a coverslip and re-immersed in distilled water. Thus the mountant was aqueous, as in the biological imaging of the current study. The beads were imaged, using an inverted microscope (Eclipse Ti-E; Nikon), with a 60× oil-immersion objective lens (numerical aperture 1.40, Plan Apo VC; Nikon), with a 0.7× intermediate coupler, and with an exposure time of 10 ms. The beads were excited by 490-nm LED at 1% intensity. A filter cube had the properties: 490/20-nm EX, 510-nm DCLP, 520-nm long-pass EM. The images were acquired with the EMCCD camera (iXon 860; Andor Technology) in a full-frame format without binning, an EM gain of 100, 14-bit 10-MHz readout, in an internal and kinetic mode (with frame transfer). The voxel size was 0.573 μm × 0.573 μm × 0.3 μm in the x-, y- and z-directions. Z-axis height was controlled manually, using the numerical distance readout of the Perfect Focus System (Nikon). In addition to the plane focused on the beads (height of 0 μm), 20 sections above and 20 sections below were imaged, with a total of 41 sections, i.e. a total height of 41 × 0.3 μm = 12.3 μm. At each height, 100 image frames were averaged, using the average-Z projection function of ImageJ (“Image/Stacks/Z project”) after the background image defined as the shutter-closed image had been subtracted. Side-view images were prepared, using an orthogonal view function of ImageJ (“Image/Stacks/Orthogonal Views”).

An intensity histogram of the beads was generated by assigning regions of interest (ROI’s, 5 × 5 pixels) to individual fluorescent puncta. For each ROI, average intensity of the pixels was used to represent the ROI intensity. The histogram was fit using a Gaussian function: N(I) = A × exp[-(I - μ)^2^ / (2 × σ^2^)], where I is the intensity in an arbitrary unit, N(I) is the number of beads at a specified intensity, A is the peak number, μ is the average intensity, and σ is the standard deviation.

Spatial resolution was defined as a width of the PSF at 50% peak height (full width at half maximum, FWHM). Assuming an elliptical PSF, two-dimensional Gaussian function was used to fit the PSF in the x-y plane at height 0 μm. The Gaussian function was expressed as: I = I_0_ + A × exp{[-(x - μ_x_)^2^ / (2 × σ_x_^2^)] + [-(y - μ_y_)^2^ / (2 × σ_y_^2^)]}, where I is the intensity in an arbitrary unit, I_0_ is the background intensity, A is the peak intensity, x and y are x- and y-coordinates, μ_x_ and μ_x_ are the average intensities along x- and y-axes, and σ_x_ and σ_y_ are the standard deviations along x- and y-axes. After averaging the standard deviations of the Gaussian function [σ = (σ_x_ + σ_y_) / 2], FWHM was calculated as: 2 × sqrt(2 × ln2) × σ = 2.35482 × σ.

## Supplementary Figures

**Fig. S1.**
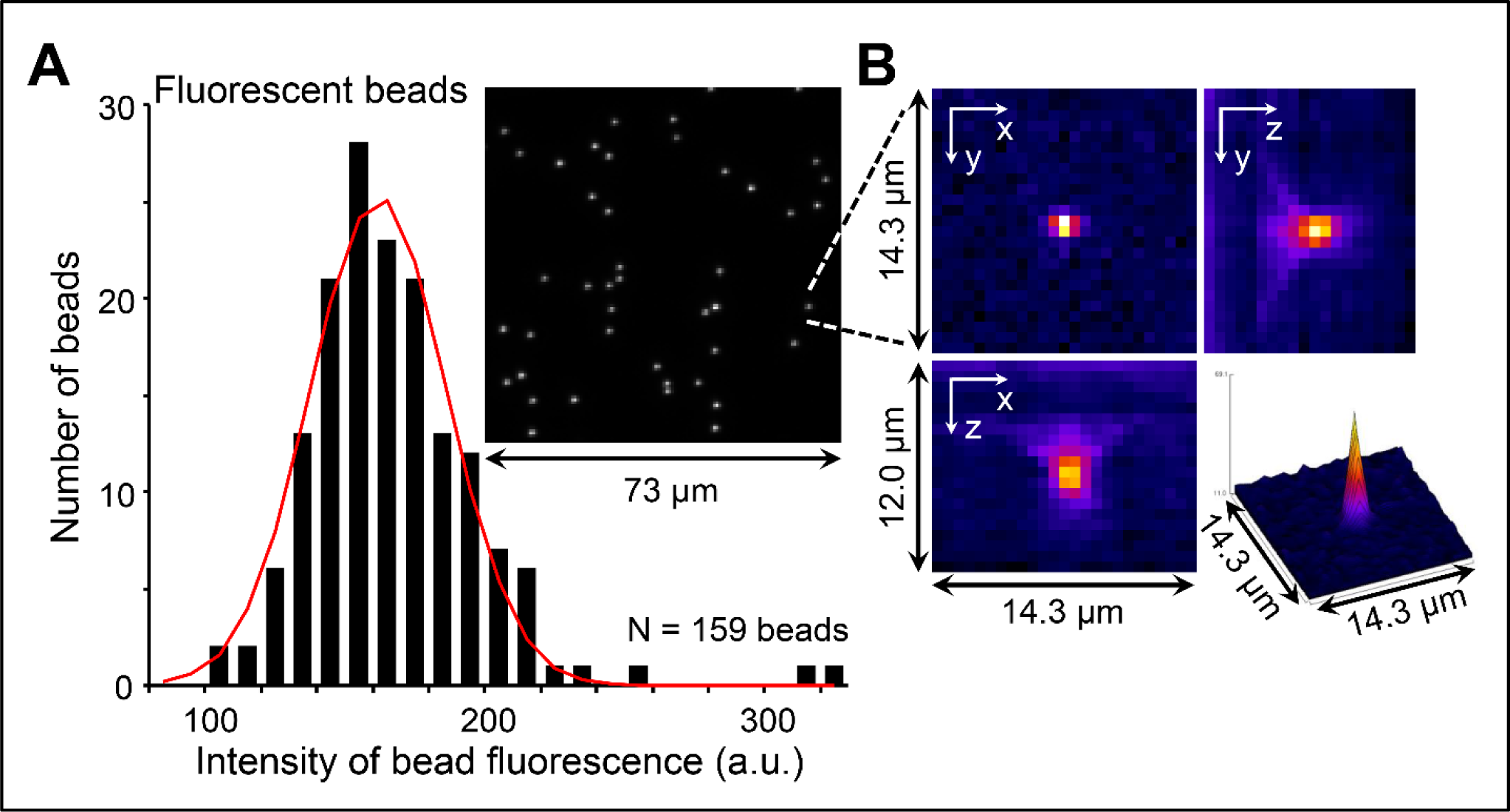
Point-spread function (PSF) analyzed with sub-resolution fluorescent beads. (**A**) Intensity histogram of punctate signals (159 signals out of 4 coverslips). The histogram was fit using a Gaussian distribution, with the peak height of 26.03, average intensity of 156.49 (arbitrary units, a.u.), and standard deviation of 23.61 (a.u.). A single Gaussian fit indicates that the signals were made up of individual fluorescent beads, i.e. beads without aggregations. (**B**) Three-dimensional PSF of a single point source. The x- and y-axes determine the ordinary imaging plane in parallel with the microscope stage, and the z-axis determines the direction of the optical path, which is perpendicular to the x-y plane. Three panels with x-, y- and z-axis notations show the PSF projected to a plane specified by the pair of axes. The panel at bottom right shows a surface plot of a bead in focus (i.e. at the z-height of 0 μm), using a color lookup table of “Fire”.

**Fig. S2.**
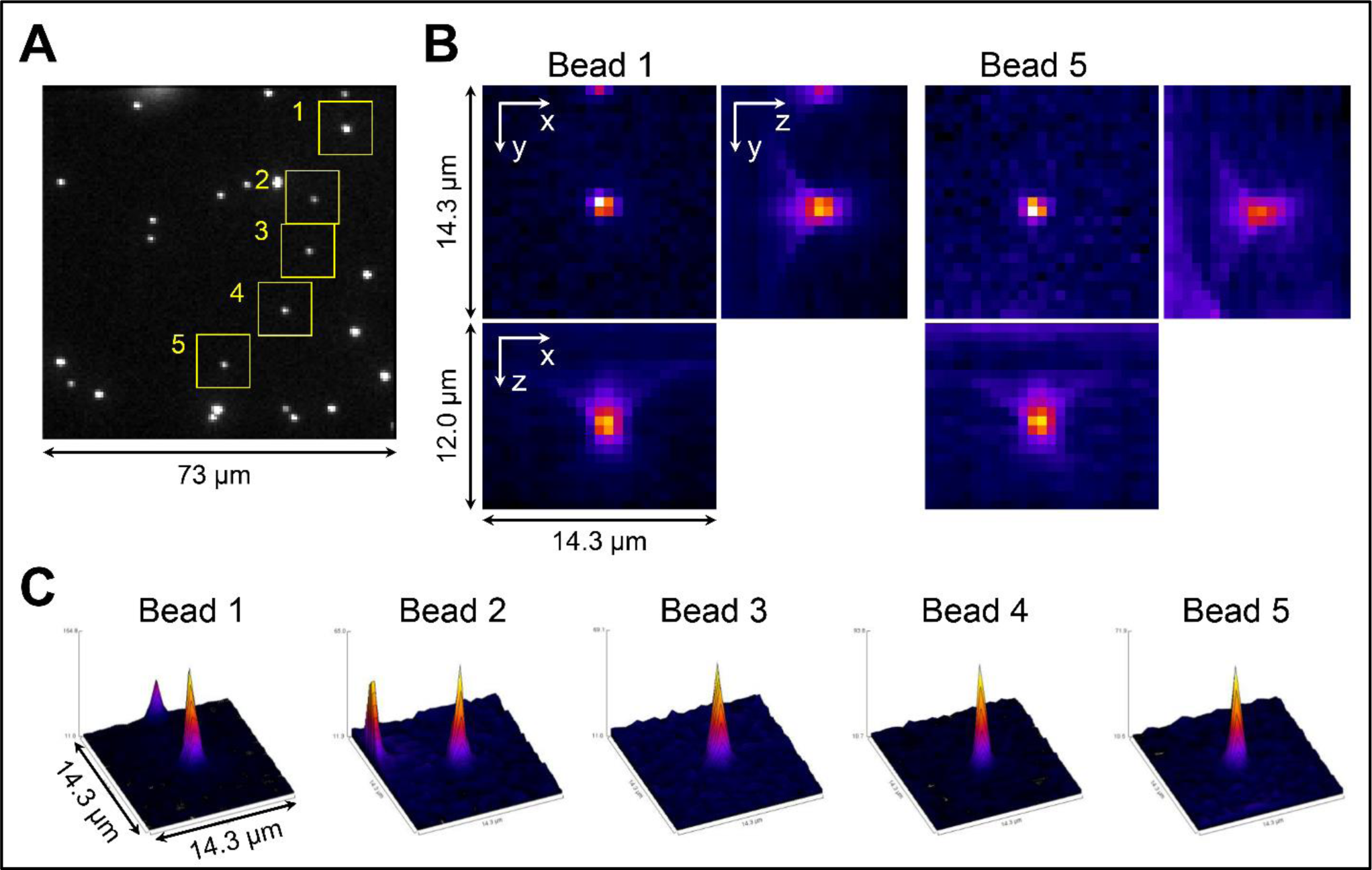
Spatial resolution, calculated based on PSF. (**A**) Image field used to measure the spatial resolution. Yellow squares indicate five regions for the measurements. (**B**) Two representative three-dimensional PSF’s. (**C**) Surface plots of the PSF, with the normalized peak intensities. The averaged standard deviation of PSF Gaussian fit of was 0.263 ± 0.004 μm (mean ± sem), and the full width at half maximum (FWHM) was 0.618 ± 0.008 μm (mean ± sem, n = 5 beads).

